# FIXATION MATTERS: MULTIDIMENSIONAL EFFECTS OF CHEMICAL PREPARATION ON MORPHOLOGY AND SURFACE TEXTURE OF EXTREMOPHILIC BACTERIA

**DOI:** 10.64898/2026.06.14.732170

**Authors:** Fátima Silvina Galván, Virginia Helena Albarracín

## Abstract

Scanning electron microscopy (SEM) is widely used to investigate bacterial surface architecture; however, sample preparation protocols may introduce structural artefacts that compromise the interpretation of morphometric and ultrastructural features. This issue becomes especially relevant for extremophilic microorganisms, whose specialized cell envelopes may respond differently to chemical fixation. In this study, we evaluated the effects of two aldehyde-based fixation protocols, 2.5% glutaraldehyde and Karnovsky’s solution, combined with different fixation times (1, 3, and 24 h) and the presence or absence of osmium tetroxide (OsO_4_) post-fixation, on three Gram-positive bacterial strains: the polyextremophiles *Exiguobacterium* sp. S17 and *Nesterenkonia* sp. Act20, and the mesophile *Kocuria rosea* CH-021. Morphological preservation was assessed using morphometric parameters, including cellular area and surface-to-volume ratio, together with texture analysis based on Haralick descriptors derived from grey-level co-occurrence matrices (GLCM). Results showed that fixation conditions significantly affected morphometric and textural features in a strain-dependent manner. Although overall morphology appeared preserved, quantitative analyses revealed marked differences in surface texture and structural integrity among treatments. Osmium tetroxide modified morphometric and textural parameters, although its effects varied among strains and fixation conditions. These findings highlight the importance of integrated quantitative approaches combining cellular geometry and surface texture analysis for the evaluation of SEM preparation protocols in structurally specialized microorganisms.

## 1. INTRODUCTION

Bacterial cell morphology is a fundamental physiological trait closely associated with growth, nutrient uptake, motility, surface colonization, and environmental adaptation (Chao & Zhang, 2011; Ojkic et al., 2019; Zhu et al., 2021). These morphological features are largely determined by the architecture and mechanical organization of the bacterial cell envelope.

The bacterial cell wall, primarily composed of covalently cross-linked peptidoglycan (PG), functions as a dynamic exoskeleton that withstands intracellular turgor pressure while allowing controlled expansion during growth and division. In Gram-positive bacteria, the thick PG layer is associated with teichoic and lipoteichoic acids, which contribute to envelope rigidity, surface charge, and selective permeability (Elbaz & Ben-Yehuda, 2010; Zhu et al., 2021). Beyond structural support, the cell envelope also participates in cell adhesion, intercellular interactions, and environmental sensing (Touhami et al., 2004).

Extremophilic and polyextremophilic bacteria exhibit specialized adaptations in their cell envelopes that enable survival under conditions such as high salinity, extreme pH, radiation exposure, and osmotic stress. These adaptations involve modifications in membrane composition, peptidoglycan organization, transport systems, and surface-associated structures that contribute to cellular stability under environmental stress (Rodrigues et al., 2008; Belfiore et al., 2013; Zannier et al., 2019). Among Gram-positive extremophiles, species of the genera *Exiguobacterium* and *Nesterenkonia* have been widely used as models for studying cellular responses to environmental stress and structural adaptation of the bacterial envelope (Aliyu et al., 2016; Zannier et al., 2022; Galván et al., 2025). Their structurally specialized envelopes make these organisms particularly relevant for evaluating how fixation and sample preparation protocols influence ultrastructural preservation in scanning electron microscopy (SEM).

SEM has become a central tool for investigating bacterial surface architecture due to its high spatial resolution, large depth of field, and ability to reveal three-dimensional surface topography (Kim, 2016; Zhang et al., 2017). In extremophilic and environmentally adapted bacteria, SEM has been widely used to examine structural features associated with cell envelope organization and stress-related surface responses (Albarracín et al., 2010; Benimeli et al., 2011; Bequer Urbano et al., 2013). These studies highlight the value of SEM for correlating surface morphology with physiological adaptation and ultrastructural organization.

Traditional SEM requires extensive sample preparation, including chemical fixation, dehydration, critical point drying, and conductive coating. Chemical fixation aims to stabilize cellular structures and minimize autolysis through biomolecular crosslinking. Aldehyde-based fixatives, such as paraformaldehyde and glutaraldehyde, stabilize cellular structures through protein crosslinking reactions (Kiernan, 2000; Park et al., 2016), while post-fixation with osmium tetroxide enhances membrane stabilization and contrast through oxidative reactions with unsaturated lipids (Peracchia & Mittler, 1972; Sorrivas de Lozano et al., 2014; Czerwińska-Główka & Krukiewicz, 2021).

Karnovsky’s solution, which combines paraformaldehyde and glutaraldehyde in buffered solution, is widely used to exploit rapid penetration by formaldehyde and stronger crosslinking by glutaraldehyde (Morris, 1965; Kiernan, 2000). Despite these established protocols, chemical fixation may introduce structural alterations such as shrinkage, membrane distortion, or extraction of soluble components, particularly during dehydration (Liu et al., 2012; Singh et al., 2019). Although cryogenic approaches reduce such artifacts, their technical and economic constraints limit widespread implementation (Touhami et al., 2004; Talbot and White, 2013; Idziak et al., 2023).

Importantly, although chemical fixation protocols are widely used to preserve bacterial morphology, their quantitative effect on morphometric and surface texture parameters has rarely been systematically evaluated, particularly in extremophilic Gram-positive bacteria. Most assessments rely on qualitative visual inspection, which introduces subjectivity and limits reproducibility.

To address this gap, the present study investigates the effects of two commonly used aldehyde-based fixation protocols (2.5% glutaraldehyde and Karnovsky’s solution), applied at different fixation times and with or without osmium tetroxide post-fixation, on three Gram-positive bacterial strains: the polyextremophiles *Exiguobacterium* sp. S17 and *Nesterenkonia* sp. Act20, and the mesophilic strain *Kocuria rosea* CH-021. We combine quantitative morphometric analysis (cellular area and surface-to-volume ratio) with texture analysis based on Haralick descriptors derived from gray-level co-occurrence matrices (GLCM). This integrative approach enables an objective assessment of fixation-induced variability and provides a comparative framework for evaluating SEM preparation protocols in structurally specialized bacteria.

## 2. MATERIALS AND METHODS

### 2.1. Strains and Media

All strains used in this study belong to and are curated in our *in-house* collection, *Cepario de Microorganismos Ambientales del Centro Integral de Microscopía Electrónica* (CeMAC). The collection includes the Gram-positive polyextremophilic bacterial strains *Exiguobacterium* sp. S17 and *Nesterenkonia* sp. Act20, both from the Catalogue HAAL, previously isolated from modern stromatolites and sediments surrounding Socompa Lake (3,570 m above sea level) in the Salta region of Argentina (S 24°35’34’’ W 68°12’42’’) (Ordoñez et al., 2013; Rasuk et al., 2017). In addition, the non-extremophilic Gram-positive strain *Kocuria rosea* CH-021, from the Catalogue Urban Bacteria, was originally isolated from the surface of the entrance door of the Museo Casa Histórica de la Independencia (Alonso-Reyes et al., 2025). All strains were processed under identical experimental conditions. *Exiguobacterium* sp. S17 (herein S17) and *Kocuria rosea* CH-021 (herein CH21) were cultured in Luria–Bertani (LB) medium adjusted to pH 8 and pH 7, respectively, whereas *Nesterenkonia* sp. Act20 (herein Act20) was cultivated in H medium (Galván et al., 2025). Cultures were incubated at 30°C under constant agitation (180 rpm) in their corresponding media. Cells were harvested during the exponential growth phase, when the optical density at 600 nm (OD600) reached 0.8-1.0, and were immediately processed for subsequent experiments.

### 2.2. Sample processing for Scanning Electron Microscopy

Circular glass coverslips (12 mm in diameter and 0.13 mm thick) were used as supports for scanning electron microscopy (SEM) measurements. Coverslips were initially immersed overnight in 96° ethanol and subsequently subjected to two successive 20-min washes in isopropyl alcohol and acetone. After cleaning, coverslips were rinsed with distilled water, placed in sterile Petri dishes, dried in an oven for 1 h, and sterilized with germicidal UV-C radiation for 30 min before use. A 100 µL aliquot of cell suspension was deposited onto the surface of each coverslip and allowed to adhere for 3 h at room temperature inside a laminar flow hood.

Samples were then treated with Karnovsky’s solution (a mixture of 2.66% w/v paraformaldehyde and 1.66% w/v glutaraldehyde) or 2.5% glutaraldehyde for 1, 3, or 24 h. All solutions were prepared in 0.1 M phosphate buffer (pH 7.2) to standardize experimental conditions across treatments and bacterial strains. Osmium tetroxide (OsO_4_) post-fixation was evaluated as an additional preparation step by incubating samples with 1% OsO_4_ for 1 min, whereas parallel samples were processed without osmium treatment (Figure 1).

**Figure 1.**
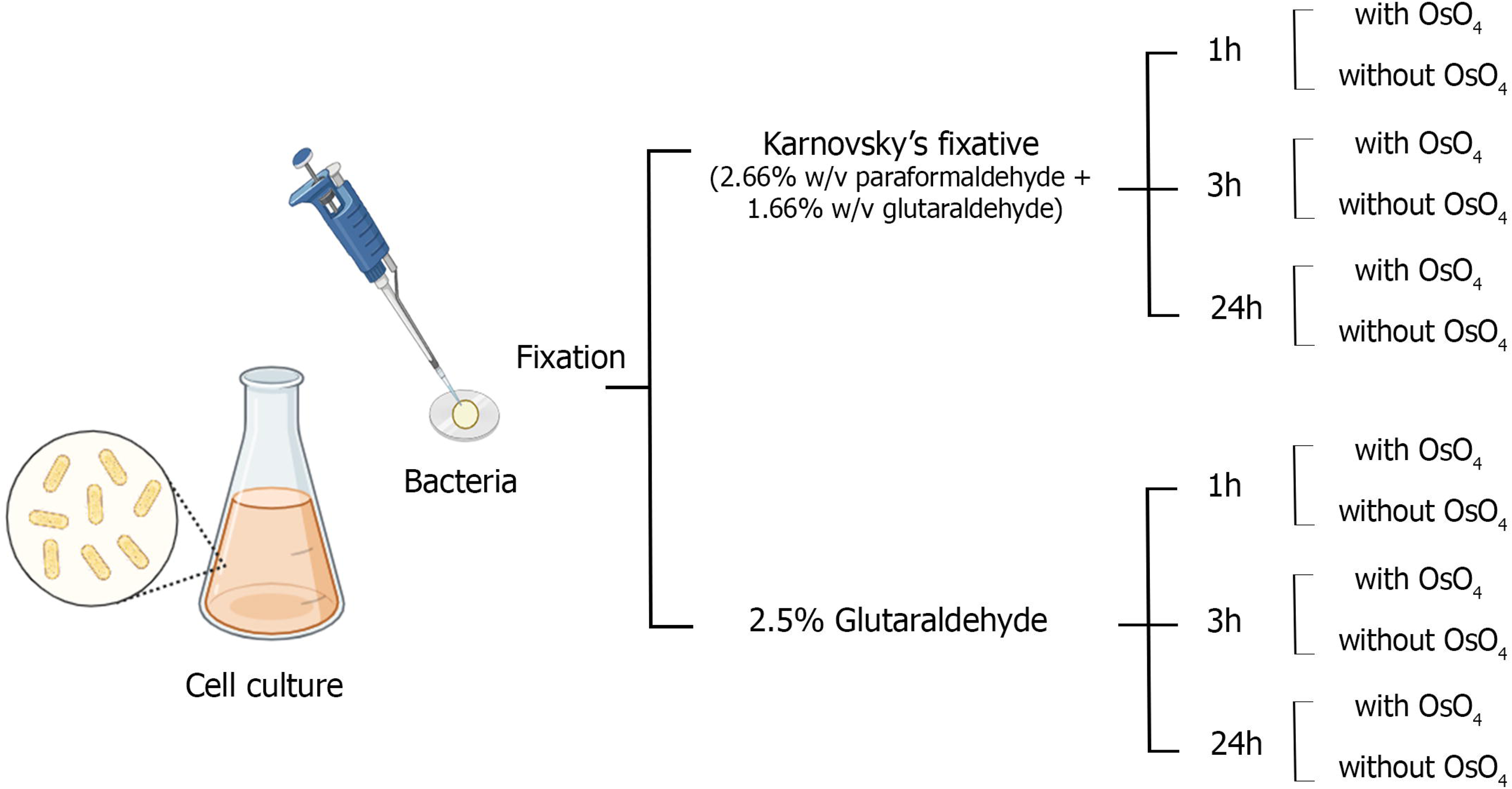
Workflow of the experimental protocols.

After fixation, samples were washed twice with 0.1 M phosphate buffer and dehydrated through a graded ethanol series (10%, 30%, 50%, 70%, 90%, and 100%), then dehydrated in 100% acetone. Critical point drying was performed using a Denton Vacuum DCP-1 apparatus. Coverslips were mounted on aluminium stubs and coated with a thin layer of gold using a JEOL JFC-1100 sputter coater. Samples were examined using a field-emission scanning electron microscope (FE-SEM, SUPRA 55VP, Zeiss, Germany) operated at 3.0 kV at the Electron Microscopy Core Facility (CIME-CONICET-UNT). Micrographs were acquired at 13,000× magnification for image analysis, while brightness and contrast settings were kept constant throughout image acquisition.

### 2.3. Morphometric analysis

The effects of the different fixation conditions on the morphology of *Exiguobacterium* sp. S17, *Nesterenkonia* sp. Act20, and *Kocuria rosea* CH-021 were evaluated by scanning electron microscopy (SEM). For comparative morphometric analysis, bacterial cells were approximated using idealized geometric models according to their predominant morphology. Rod-shaped cells of S17 were modelled as cylinders, coccobacillary cells of Act20 as cylinders with hemispherical ends, and coccoid cells of CH-021 as spheres:

- *Exiguobacterium* sp. S17:

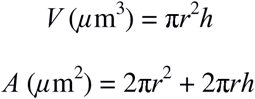
- *Nesterenkonia* sp. Act20:

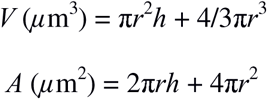
- *Kocuria rosea* CH-021:

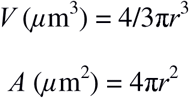

Where *r* corresponds to cell radius and *h* to total cell length, measured in micrometres (µm). Calculations were performed following established SEM-based morphometric approaches (Neumann et al., 2005; Chakravarty & Banerjee, 2008).

Cell dimensions were obtained using ImageJ version 1.54f (National Institutes of Health, USA) from micrographs acquired under each experimental condition. A total of 25 cells per condition were measured for S17 and CH-021, and 15 cells per condition for Act20 due to lower cell density. Cells undergoing division, deformation, or structural rupture were excluded from the dataset (Malyshev et al., 2024).

### 2.4. Quantitative Texture Analysis of Bacterial Images Using Haralick Features based on GLCM

For quantitative textural analysis of the bacterial surface, Haralick texture features were extracted from grey-level co-occurrence matrices (GLCM) (Haralick et al., 1973; Ríos-Díaz et al., 2009; Malyshev et al., 2024). GLCMs quantify the joint probability of pixel intensity pairs (i, j) occurring at a defined spatial relationship within the image. Matrices were computed using a pixel distance of 1 and four directions (0°, 45°, 90°, and 135°), with features averaged across orientations.

Scanning electron micrographs were acquired at 13,000× magnification with a resolution of 3072 × 2304 pixels (8-bit grayscale range: 0–255). Each analyzed micrograph corresponded to a single well-isolated bacterial cell, which was treated as an independent observational unit.

Cell selection followed predefined morphological criteria, including intactness and absence of overlap or structural damage. Selection was applied consistently across all experimental conditions.

For each cell, three 75 × 75-pixel regions of interest (ROIs) were manually defined on the cell surface to capture local textural variability. Haralick texture features, contrast (CON), correlation (COR), energy (angular second moment, ASM), entropy (ENT), and homogeneity (HOM), were computed at ROI level and averaged to yield a single representative value per cell. Thus, each cell contributed one observation per feature, and ten cells were analyzed per experimental condition.

### 2.5. Statistical Analysis

Quantitative data are presented as mean ± standard deviation (SD). Statistical analyses were performed separately for morphometric and texture datasets. For both datasets, a three-way ANOVA was used to evaluate the effects of fixative, fixation time, and post-fixation with osmium tetroxide on the measured variables. Post hoc pairwise comparisons were performed using the Tukey-Kramer test. Statistical significance was set at p < 0.05.

Morphometric parameters (cell area and surface area-to-volume ratio) and Haralick texture features were calculated at the single-cell level under standardized imaging and processing conditions. In both cases, each cell was treated as an independent observational unit. For the textural analysis, region-of-interest (ROI) measurements were nested within individual cells and averaged to obtain a single representative value per cell.

All statistical analyses, graphical representations, and data processing were performed using custom MATLAB® scripts (The MathWorks Inc., Natick, MA, USA). The original datasets and analysis scripts are provided as supplementary material (Dataset_Supplementary.xlsx and Files S1–S6).

**Figure 2.**
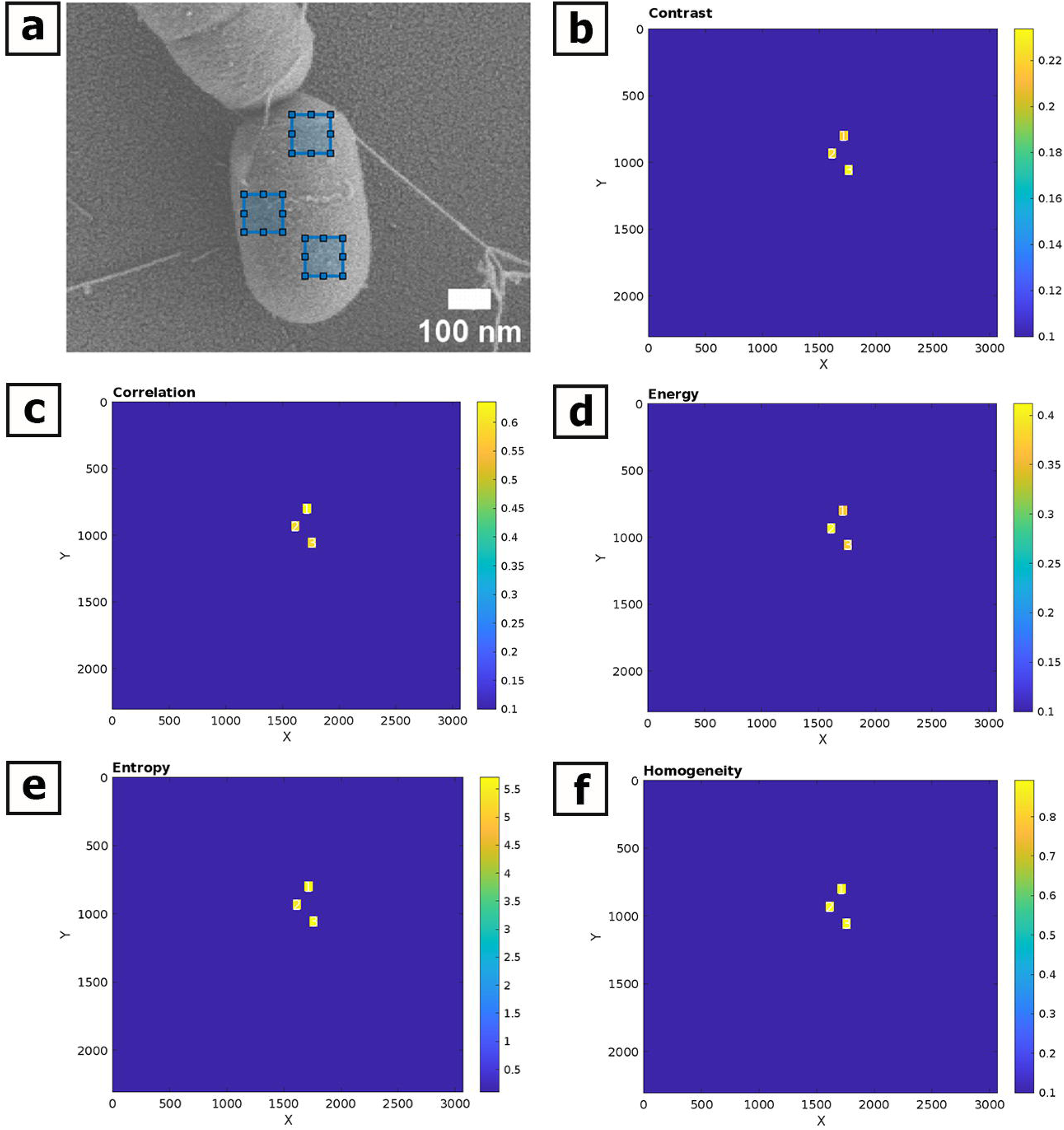
Definition of regions of interest (ROIs) used for texture analysis. Representative scanning electron micrograph showing the selection of three 75 × 75 pixel ROIs on the bacterial cell body for gray-level co-occurrence matrix (GLCM) analysis.

## 3. RESULTS

### 3.1. Fixation Protocols Induce Measurable but Strain-Dependent Morphological Variability

Cellular area and surface-to-volume ratio (S/V) were analyzed to evaluate the effects of fixation protocols on bacterial morphology (Tables 1–3; Figure 3). Distinct morphometric response patterns were observed among the three strains.

**Figure 3.**
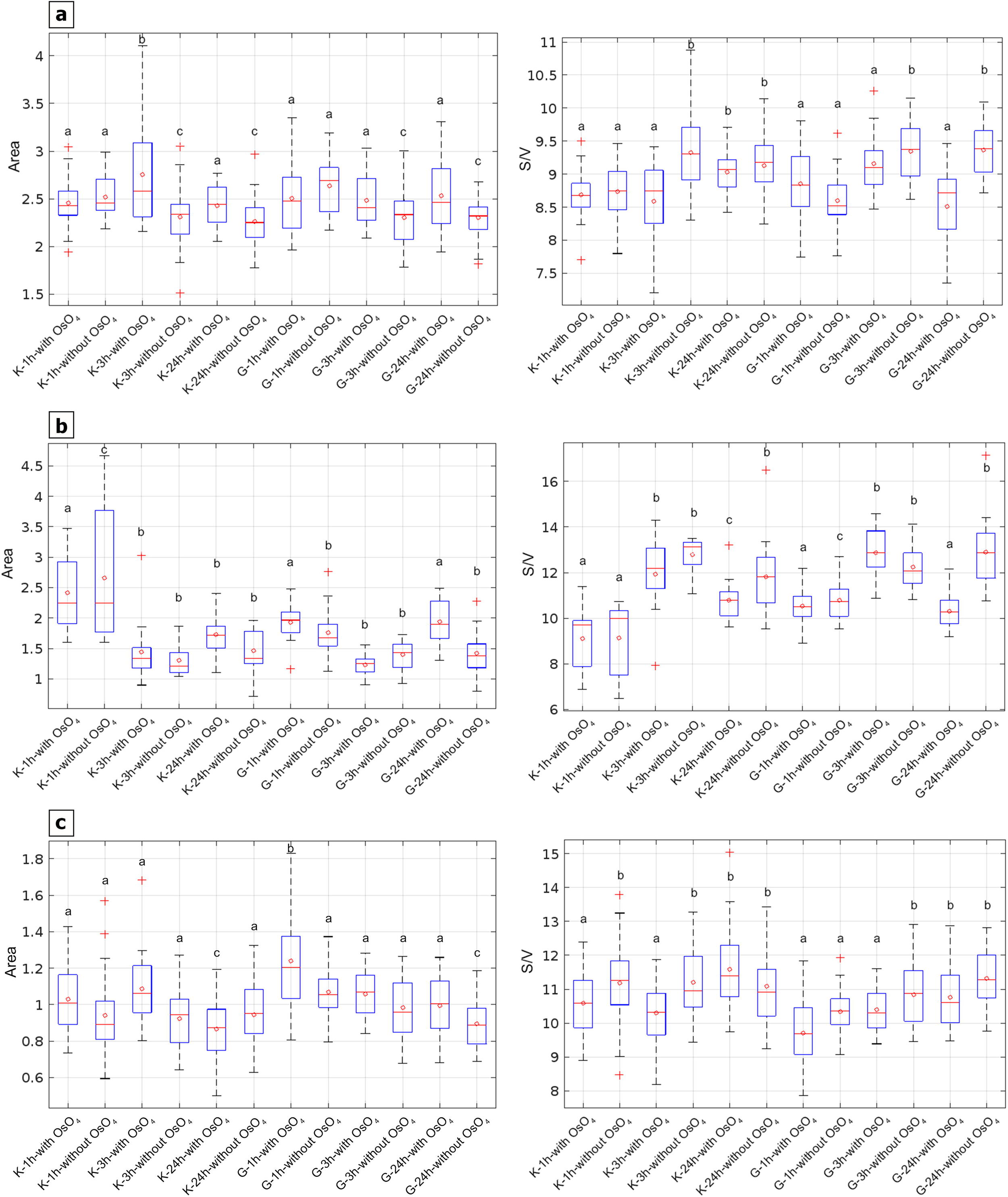
Morphometric parameters of *Exiguobacterium* sp. S17 **(a)**, *Nesterenkonia* sp. Act20 **(b)**, and *Kocuria rosea* CH-021 **(c)**, showing surface area (µm²) and surface-to-volume ratio (S/V) for each fixation treatment evaluated. Boxplots represent the distribution of individual measurements for each condition: boxes indicate the interquartile range, the central line represents the median, red open circles denote the mean value for each group, and whiskers indicate data dispersion. Different letters indicate statistically significant differences among fixation treatments based on three-way ANOVA followed by post hoc multiple comparisons (p < 0.05).

**Table 1.**
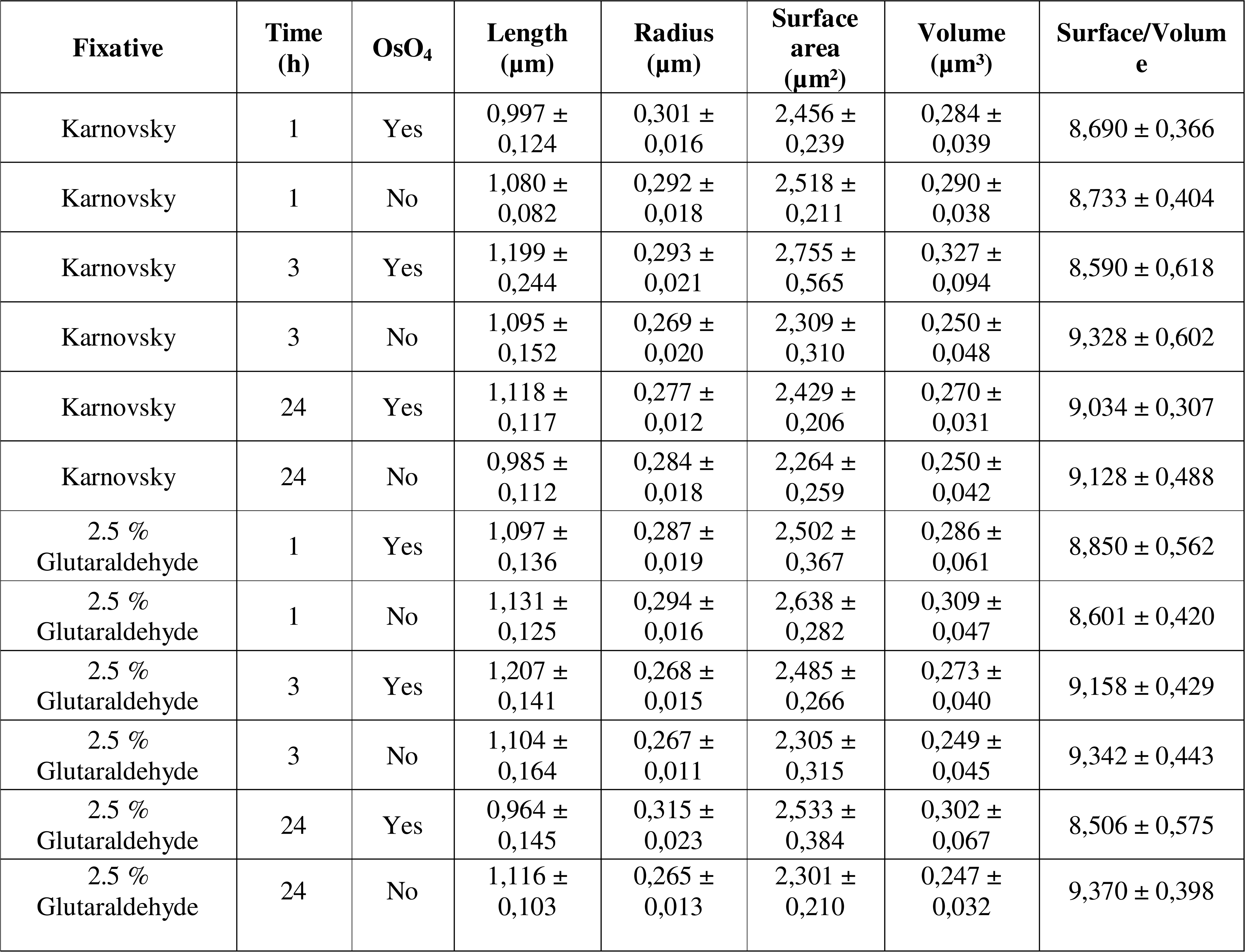
Morphometric parameters of *Exiguobacterium* sp. S17 cells, including length (µm), radius (µm), surface area (µm²), volume (µm³), and surface-to-volume ratio (S/V).

**Table 2.**
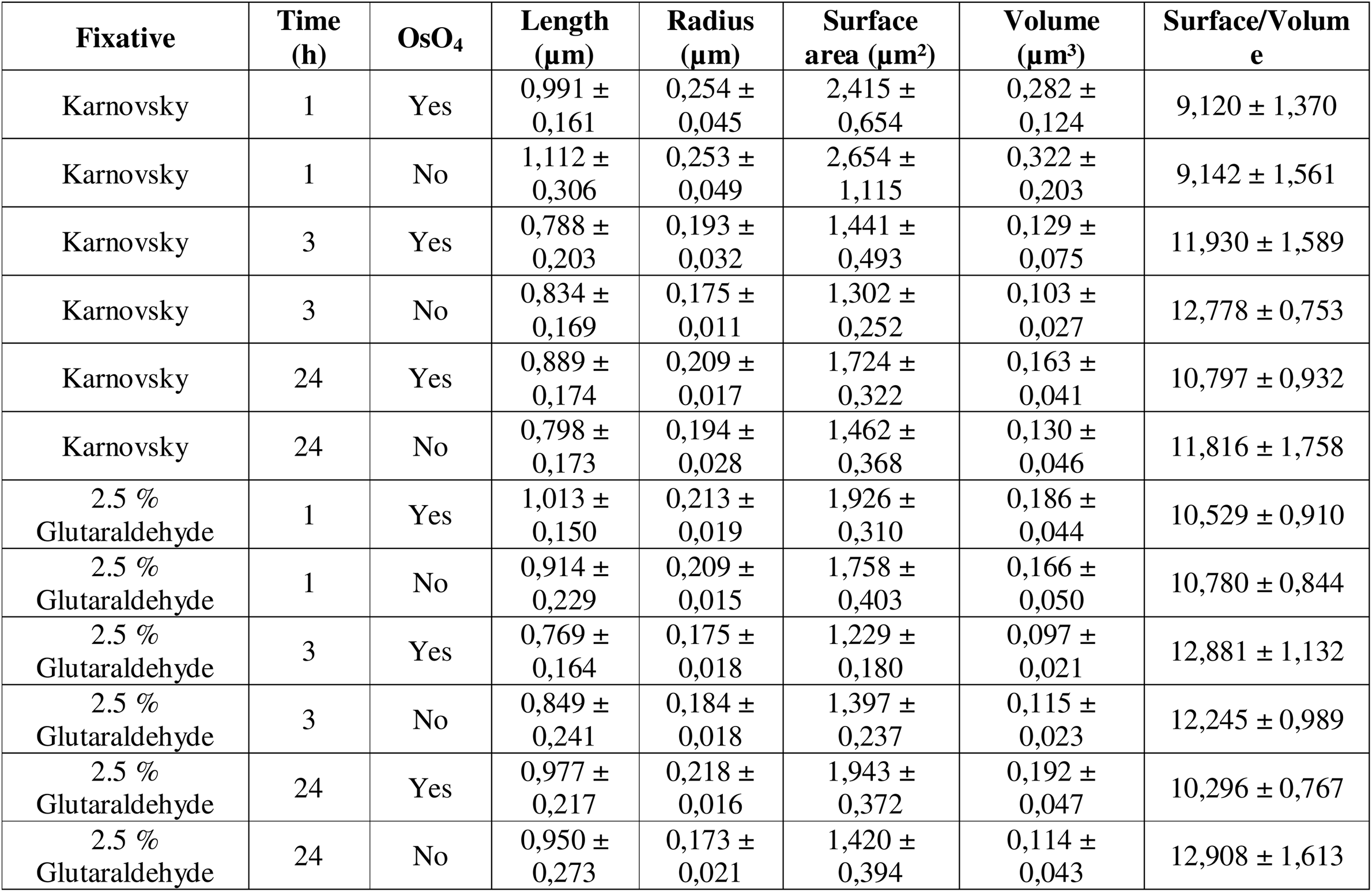
Morphometric parameters of *Nesterenkonia* sp. Act20 cells, including length (µm), radius (µm), surface area (µm²), volume (µm³), and surface-to-volume ratio (S/V).

**Table 3.**
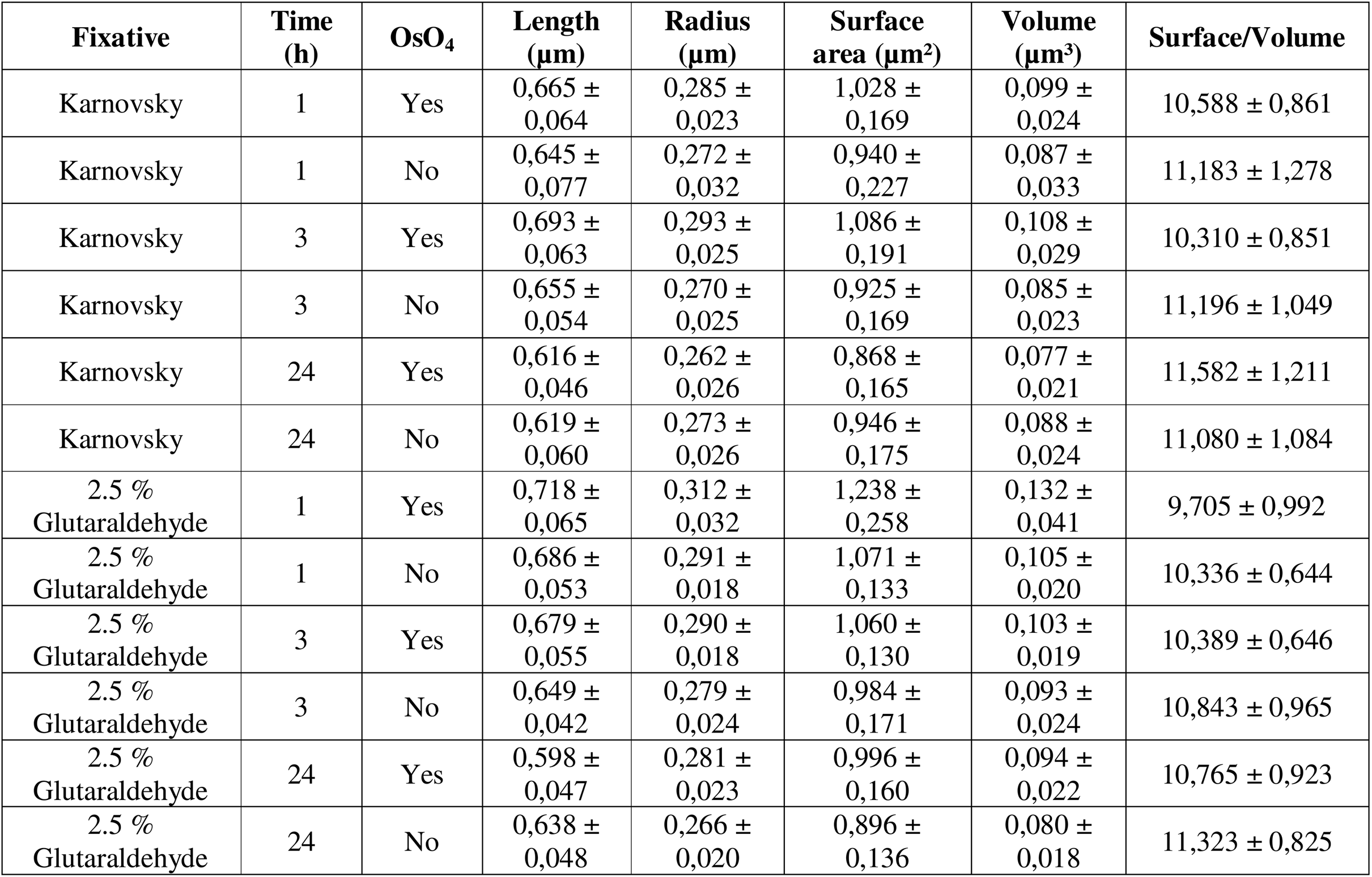
Morphometric parameters of *Kocuria rosea* CH-021 cells, including length (µm), radius (µm), surface area (µm²), volume (µm³), and surface-to-volume ratio (S/V).

S17 showed relatively stable morphometric distributions, with partial overlap among fixation treatments. In contrast, Act20 exhibited the broadest distributions of area and S/V values, indicating greater fixation-associated variability. CH-021 maintained narrow morphometric distributions across most conditions.

The standard protocol used at our institution, consisting of Karnovsky fixation for 24 h without post-fixation osmium tetroxide, yielded intermediate morphometric values across all strains. Several alternative treatments overlapped with this reference condition, whereas others differed significantly with respect to fixation time, fixative composition, and post-fixation osmium.

Together, these observations indicate that fixation effects on bacterial morphometry are strain-dependent and arise from multifactorial interactions among fixation conditions.

### 3.2. Osmium Post-Fixation Modulates Morphology Without Uniform Effects Across Strains

Osmium tetroxide post-fixation influenced bacterial morphometry differently across the three strains analyzed (Figure 3). In S17, OsO_4_-treated samples generally exhibited slightly larger cellular area values and lower S/V ratios than non-post-fixed cells, suggesting differences in the response of the cellular envelope to post-fixation.

In contrast, Act20 showed broader and partially overlapping structural profiles between post-fixed and non-post-fixed conditions, indicating greater variability in the response to osmium post-fixation. CH-021 displayed subtle shifts in area and S/V values following OsO_4_ post-fixation, although statistically distinct groups were still detected under specific fixation combinations.

Collectively, boxplot distributions and statistical groupings indicated that the effects of osmium tetroxide were strain-dependent and varied according to fixation conditions.

### 3.3. Fixation Time Contributes to Variability but Shows Species-Specific Patterns

Morphological responses varied across fixation times and differed substantially among strains.

S17 maintained relatively stable distributions of cellular area and S/V values over time, although statistically distinguishable groups were detected at specific fixation periods. In contrast, Act20 exhibited more pronounced temporal shifts in cellular geometry, indicating greater sensitivity to fixation duration. CH-021 showed limited variation throughout the evaluated time intervals, consistent with greater structural stability during prolonged fixation.

These findings indicate that fixation time contributes to bacterial morphological variability in a species-specific manner and does not follow a uniform temporal trend across fixation conditions.

### 3.4. Fixative Type Alone Does Not Determine Morphological Outcomes

Comparisons between Karnovsky solution and 2.5% glutaraldehyde revealed no consistent preservation pattern associated with either fixative alone. In S17, geometric distributions obtained with both fixatives partially overlapped across fixation times and osmium post-fixation conditions. Similar trends were observed in Act20 and CH-021, where specific fixation combinations produced distinguishable cellular responses without a uniform separation between fixatives.

The observed distributions indicate that preservation outcomes depended on the interaction among chemical composition, fixation duration, and post-fixation osmium rather than on fixative type alone.

### 3.5. Structural Preservation Emerges from Multifactorial Interactions Between Fixation Parameters

Three-way ANOVA revealed significant interaction effects among fixation parameters across the three bacterial strains (Table 4), indicating that structural preservation could not be explained by individual factors independently.

**Table 4.**
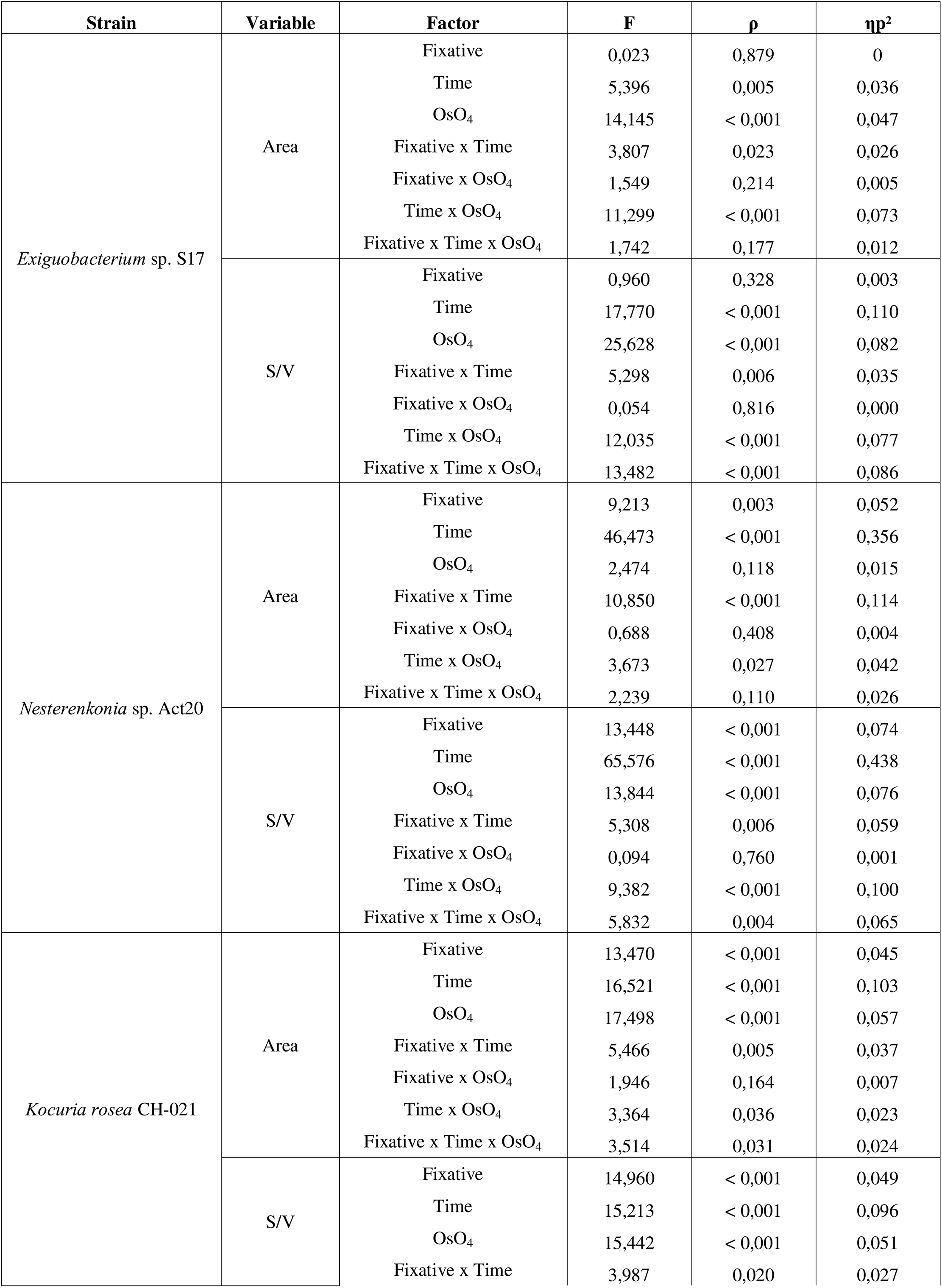
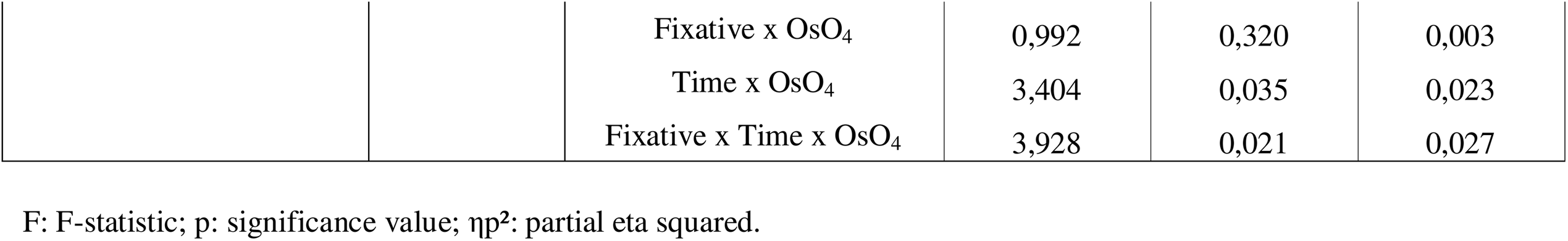
Three-way ANOVA of fixation effects on cellular area and surface-to-volume ratio in *Exiguobacterium* sp. S17, *Nesterenkonia* sp. Act20, and *Kocuria rosea* CH-021.

The relative contribution of fixation parameters differed substantially among strains. In Act20, fixation time produced the strongest effects on both cellular area and S/V ratio, reflecting marked sensitivity to fixation duration. In contrast, S17 exhibited stronger responses associated with osmium post-fixation and time-dependent interactions, whereas CH-021 displayed more moderate effect sizes distributed across multiple fixation parameters.

Several interaction terms were significant for both cellular area and S/V ratio, particularly those involving fixation time and osmium tetroxide post-fixation. These results indicate that fixation-induced structural responses were not strictly additive, but instead emerged from condition-specific relationships between fixation chemistry, fixation duration, and post-fixation treatment.

### 3.6. Surface Texture Analysis Reveals Ultrastructural Differences Not Detected by Morphometry

Haralick texture descriptors derived from grey-level co-occurrence matrices (GLCM) were used to quantify surface organization patterns in SEM micrographs (Figure 4, Tables 5–7). Across the three bacterial strains, fixation conditions associated with higher angular second moment (ASM) and homogeneity (HOM), together with lower entropy (ENT) and contrast (CON), generally corresponded to more uniform ultrastructural organization and reduced surface heterogeneity. In contrast, elevated entropy and contrast values were associated with more irregular grayscale organization and increased microstructural variability. Correlation (COR) values varied less markedly across fixation conditions than the remaining texture descriptors, indicating limited sensitivity to fixation-induced surface alterations.

**Figure 4.**
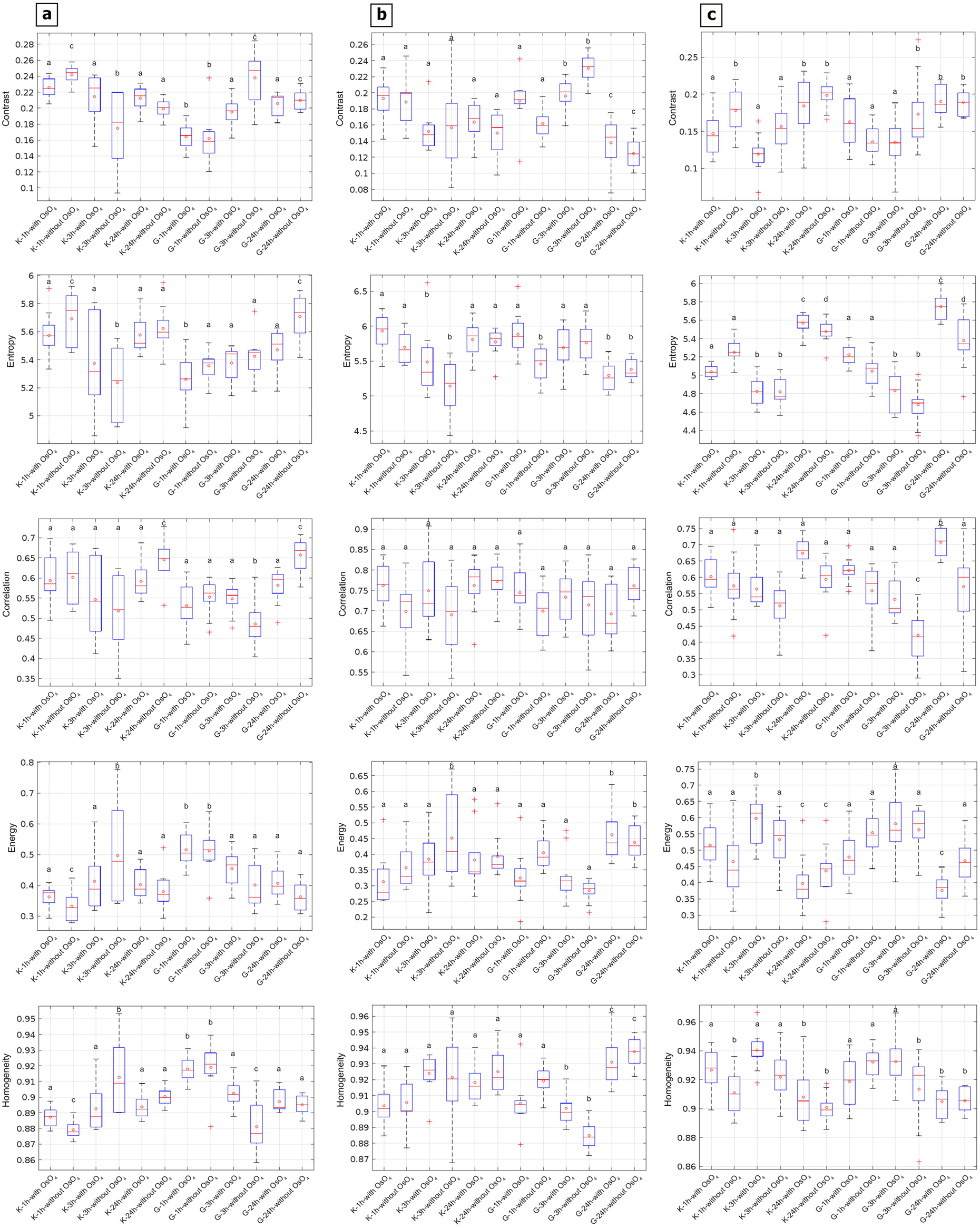
Haralick texture feature values of *Exiguobacterium* sp. S17 **(a)**, *Nesterenkonia* sp. Act20 **(b)**, and *Kocuria rosea* CH-021 **(c)**, showing contrast (CON), correlation (COR), energy (ASM), entropy (ENT), and homogeneity (HOM), calculated from gray-level co-occurrence matrices (GLCM) over regions of interest in SEM images for each fixation treatment evaluated. Boxplots represent the distribution of individual measurements for each condition: boxes indicate the interquartile range, the central line represents the median, red open circles denote the mean value for each group, and whiskers indicate data dispersion. Different letters indicate statistically significant differences among fixation treatments based on three-way ANOVA followed by post hoc multiple comparisons (p < 0.05).

**Table 5.**
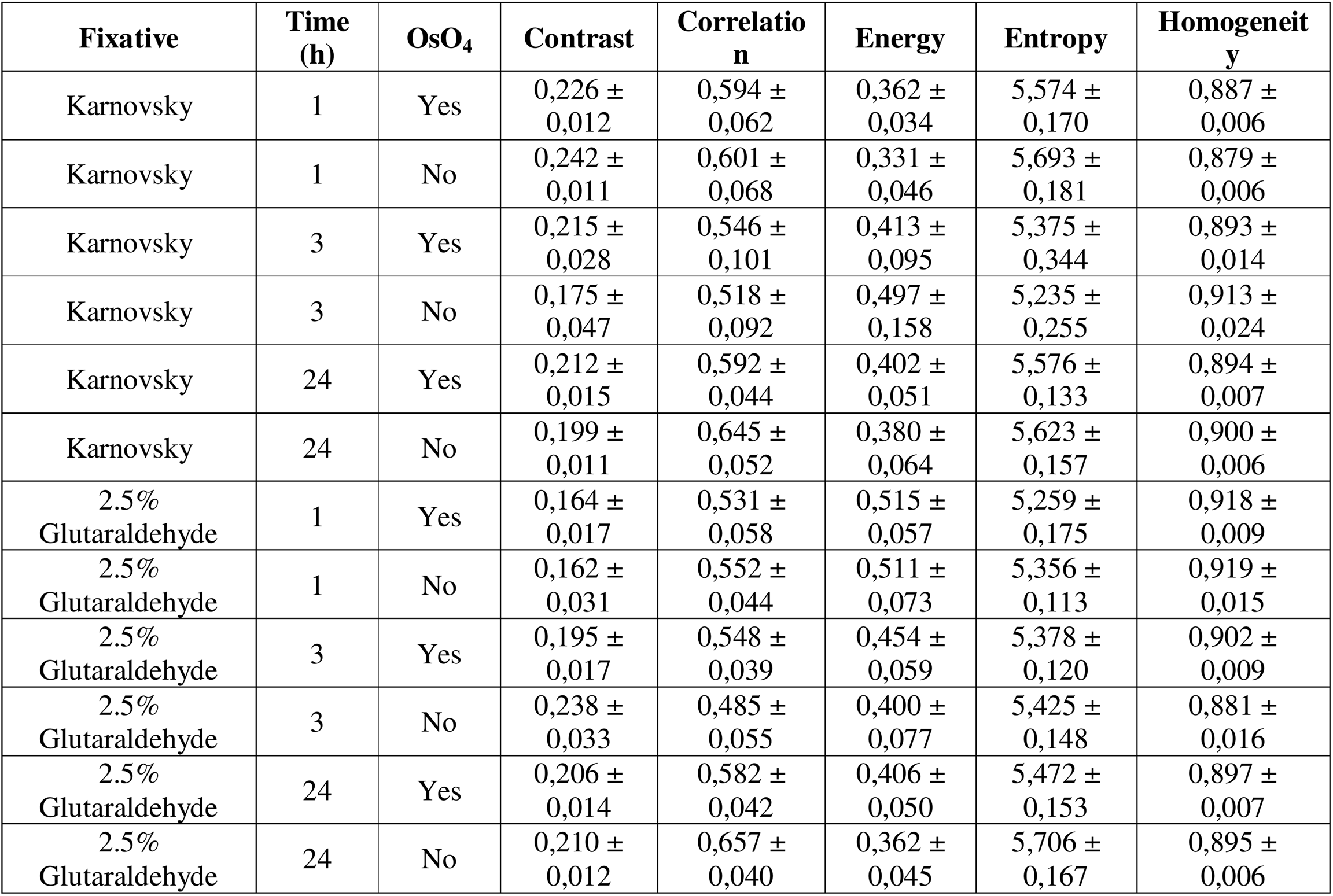
Haralick textural parameters of *Exiguobacterium* sp. S17 cells, including contrast (CON), correlation (COR), energy (ASM), entropy (ENT), and homogeneity (HOM), calculated from gray-level co-occurrence matrices (GLCM) over regions of interest in SEM images.

**Table 6.**
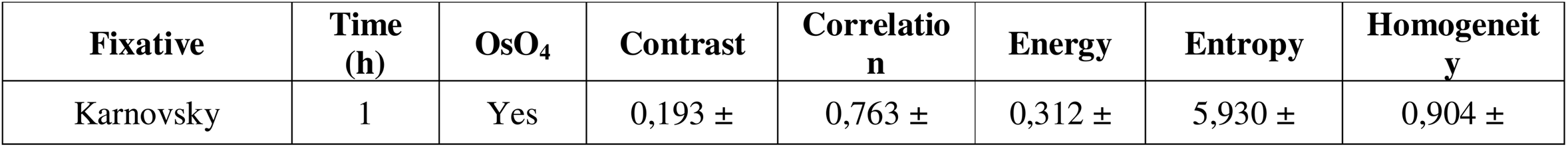
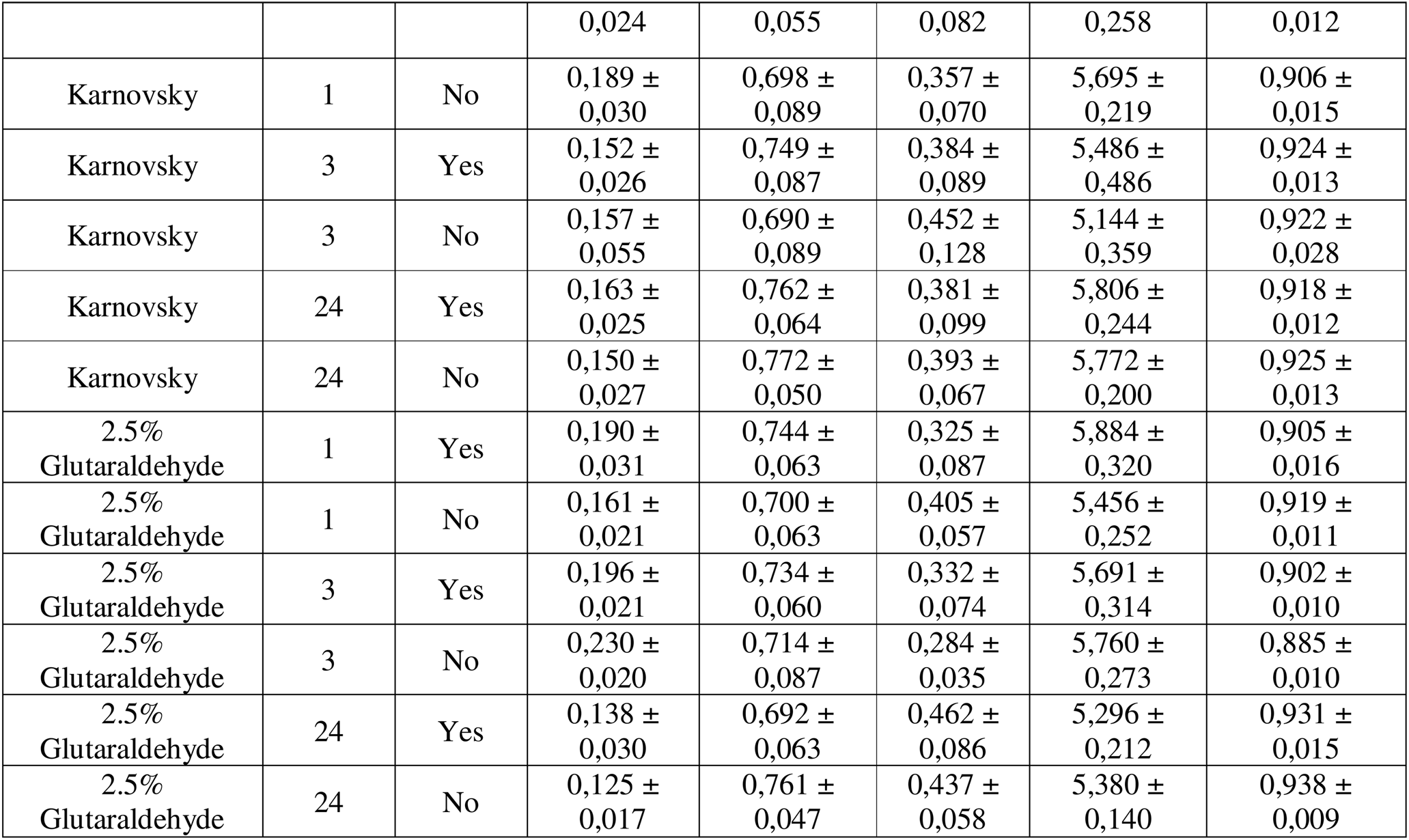
Haralick textural parameters of *Nesterenkonia* sp. Act20 cells, including contrast (CON), correlation (COR), energy (ASM), entropy (ENT), and homogeneity (HOM), calculated from gray-level co-occurrence matrices (GLCM) over regions of interest in SEM images.

**Table 7.**
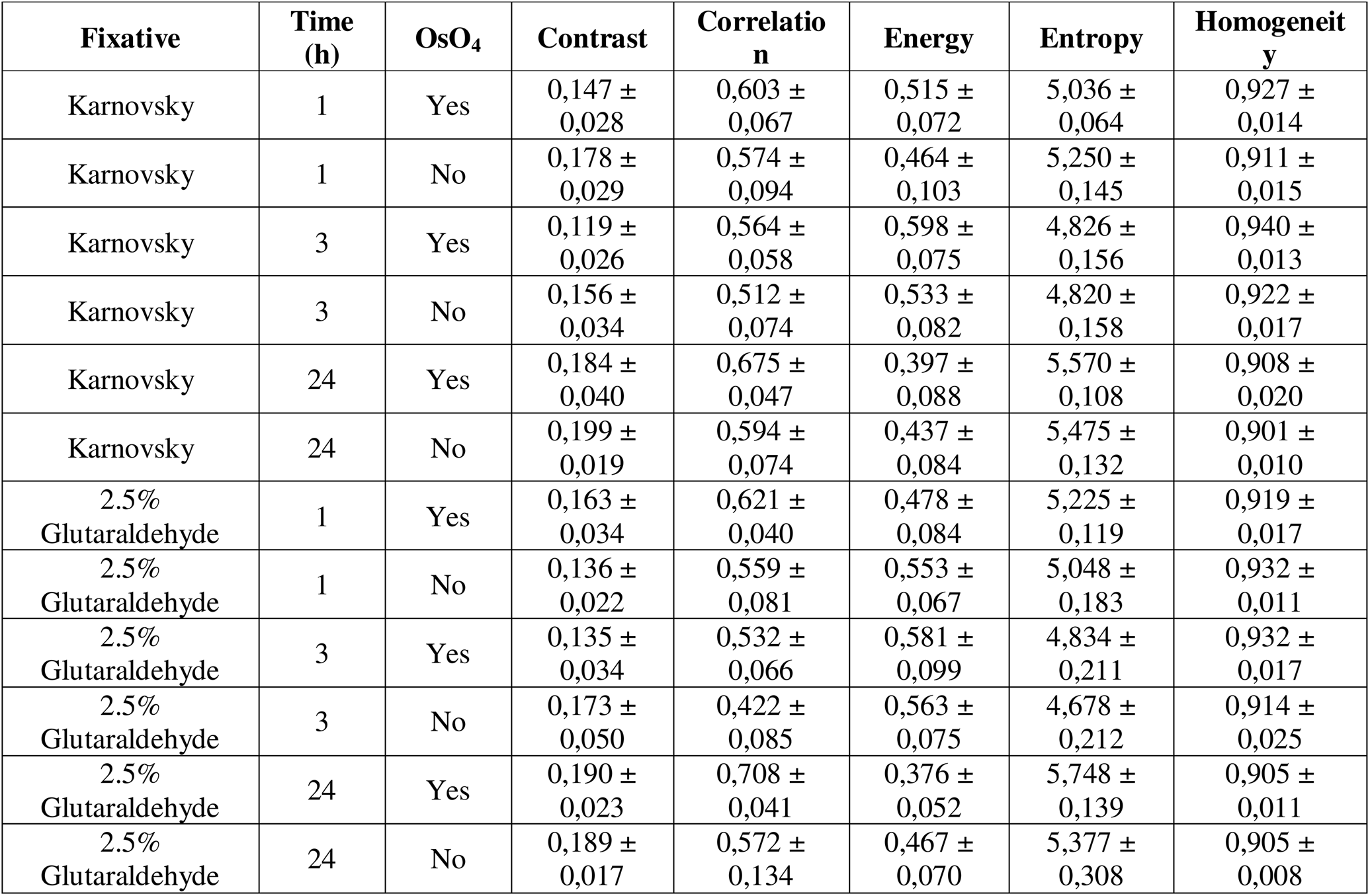
Haralick textural parameters of *Kocuria rosea* CH-021 cells, including contrast (CON), correlation (COR), energy (ASM), entropy (ENT), and homogeneity (HOM), calculated from gray-level co-occurrence matrices (GLCM) over regions of interest in SEM images.

In S17, short fixation with 2.5% glutaraldehyde, with or without OsO_4_ post-fixation, produced texture profiles with increased ASM and HOM values together with reduced contrast, consistent with relatively homogeneous surface patterning. In Act20, prolonged fixation combined with osmium post-fixation was associated with more homogeneous ultrastructural organization than several alternative treatments. Nevertheless, texture descriptors across the full set of fixation conditions remained broadly dispersed, indicating persistent sensitivity of surface texture to fixation parameters. In CH-021, intermediate fixation periods combined with osmium post-fixation produced more homogeneous ultrastructural organization relative to several alternative treatments.

Importantly, several fixation conditions that produced comparable cellular area and S/V profiles exhibited markedly different surface texture patterns. These findings indicate that dimensional preservation alone does not necessarily reflect equivalent conservation of surface microstructure.

### 3.7. Fixation-Dependent Differences in Ultrastructural Surface Features

SEM micrographs revealed variations in ultrastructural surface features across fixation conditions in the three bacterial strains analyzed (Figures 5–7). Differences were observed in the appearance of tubular structures, membrane protrusions, extracellular material, and surface irregularities.

**Figure 5.**
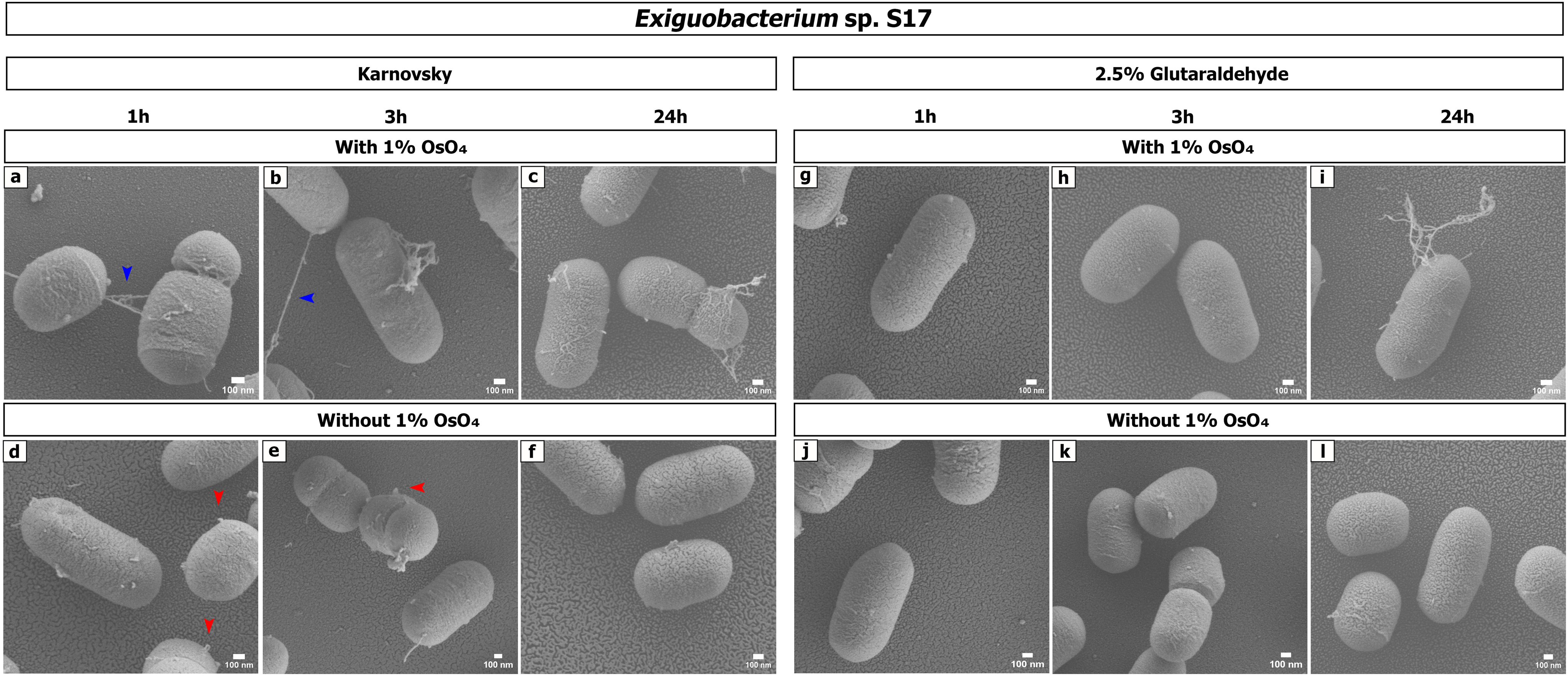
Scanning electron microscopy (SEM) images of *Exiguobacterium* sp. S17 cells subjected to different fixation treatments. Representative micrographs illustrate the effects of fixative type, fixation time, and treatment with or without OsO_4_ post-fixation on cell surface morphology and structural preservation. Blue arrows indicate tubular structures, whereas red arrows indicate membrane protrusions. Scale bars: 100 nm.

**Figure 6.**
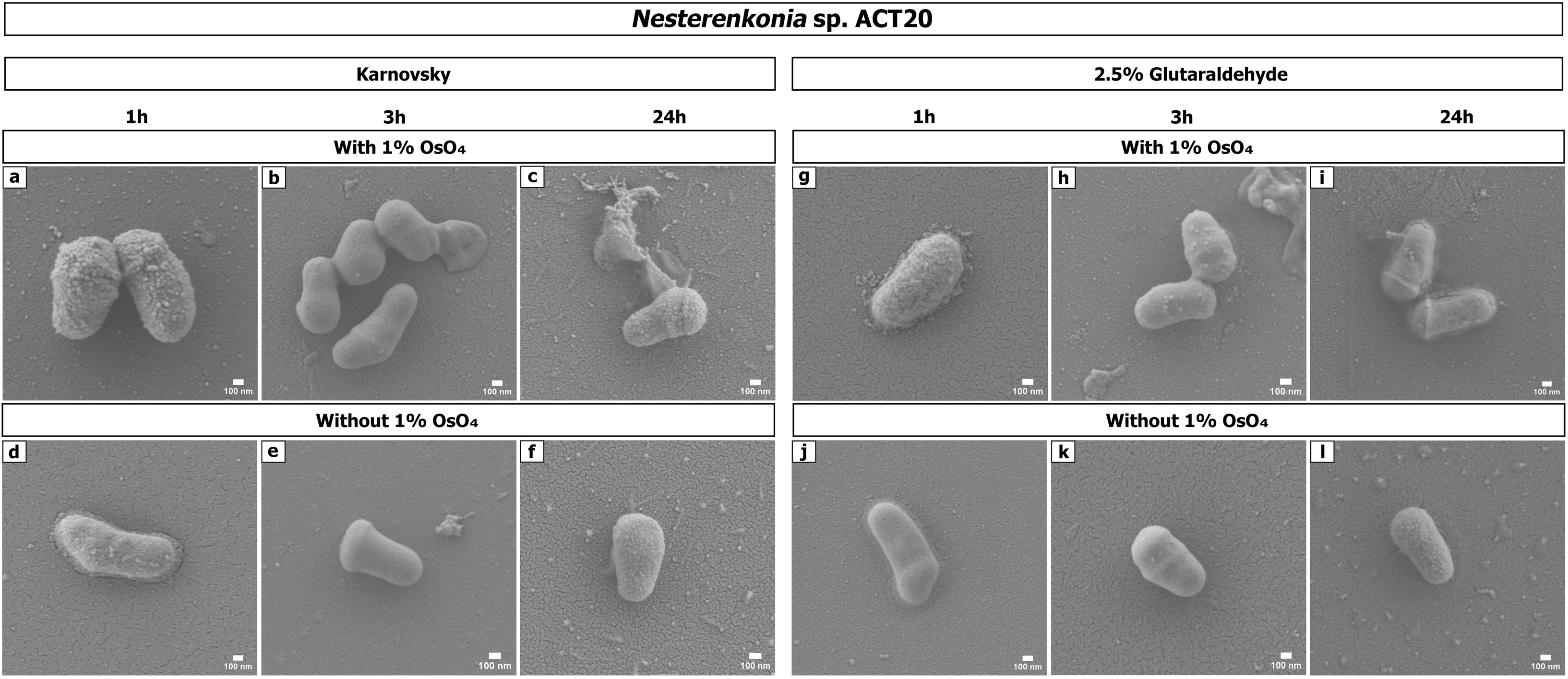
Scanning electron microscopy (SEM) images of *Nesterenkonia* sp. Act20 cells subjected to different fixation treatments. Representative micrographs illustrate the effects of fixative type, fixation time, and treatment with or without OsO_4_ post-fixation on cell surface morphology and structural preservation. White arrows indicate extracellular material surrounding the cells. Scale bars are 100 nm.

**Figure 7.**
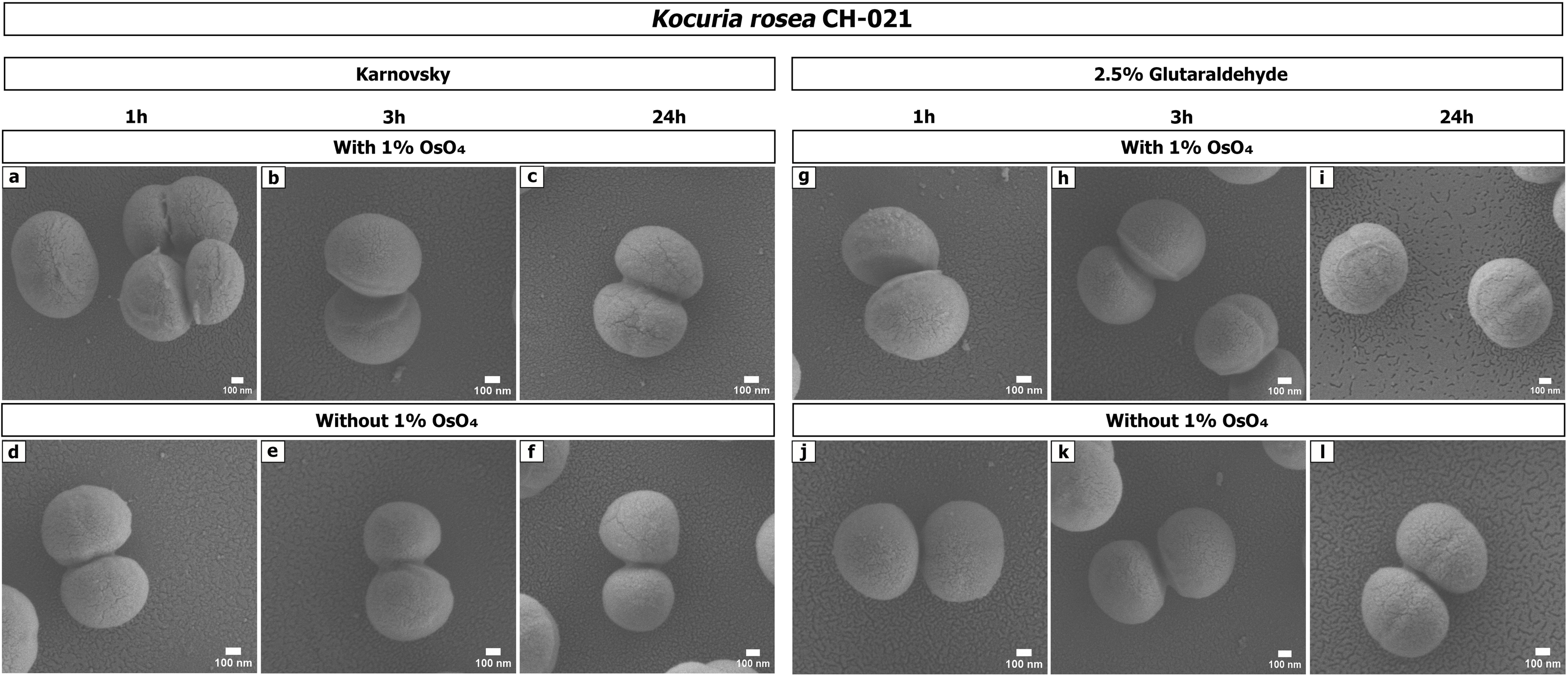
Scanning electron microscopy (SEM) images of *Kocuria rosea* CH-021 cells subjected to different fixation treatments. Representative micrographs illustrate the effects of fixative type, fixation time, and treatment with or without OsO_4_ post-fixation on cell surface morphology and structural preservation. Scale bars are 100 nm.

In S17, filamentous intercellular structures morphologically resembling previously described nanotube-like formations were detected predominantly under Karnovsky fixation combined with osmium tetroxide post-fixation, with apparent diameters ranging from approximately 20–30 nm (Figure 5, blue arrow). Under fixation conditions without osmium post-fixation, nanoscale membrane protrusions and irregular surface extensions were observed more frequently (Figure 5, red arrow).

Extracellular material surrounding bacterial cells was observed in both S17 and Act20. In Act20, accumulation of extracellular material was particularly evident following fixation with 2.5% glutaraldehyde, occasionally reducing the visual definition of cell boundaries (Figure 6, white arrow). Vesicle-like surface protrusions were also detected under multiple fixation conditions, particularly in samples subjected to osmium tetroxide post-fixation. In addition, structures compatible with cellular debris or localized material redistribution were detected under specific preparation conditions.

Taken together, these observations indicate that the visualization of ultrastructural surface features varied according to fixation and post-fixation conditions, suggesting that SEM-based structural interpretation may depend partly on sample preparation protocols. Because all images were obtained after chemical fixation, dehydration, critical-point drying and metal coating, the contribution of preparation-induced artifacts cannot be completely excluded.

## 4. DISCUSSION

In the present study, we demonstrate that fixation protocols induce measurable but strongly strain-dependent effects on both morphometric and textural properties of extremophilic Gram-positive bacteria in scanning electron microscopy (SEM). While fixation-induced artifacts have been widely reported in microorganisms (Chao & Zhang, 2011), their quantitative variability in extremophiles remains insufficiently characterized. Our results reveal that global morphology and surface texture may respond differently to fixation conditions, highlighting the need for integrated structural assessment.

Cellular morphology in SEM reflects the combined effects of chemical fixation and subsequent sample processing steps. Aldehyde-based fixatives, such as paraformaldehyde and glutaraldehyde, primarily stabilize proteins through cross-linking reactions, whereas osmium tetroxide enhances membrane stabilization and surface contrast through oxidative reactions with unsaturated fatty acids (Kiernan, 2000; Czerwińska-Główka & Krukiewicz, 2021). However, subsequent dehydration and critical point drying steps may still induce structural collapse or shrinkage when chemical stabilization is incomplete (Gusnard & Kirschner, 1977). Accordingly, SEM-derived morphometric measurements reflect the combined influence of intrinsic cellular architecture and preparation-induced structural modifications.

Within this framework, cellular area and surface-to-volume ratio (S/V) provide complementary indicators of structural preservation. Area reflects global dimensional changes in cell morphology, whereas S/V integrates geometric and volumetric relationships associated with diffusion constraints and envelope integrity (Harris & Theriot, 2018; Ojkic et al., 2019). Because S/V is mathematically coupled to cellular volume under approximate shape conservation, it is particularly sensitive to subtle fixation-induced deformation that may not be visually apparent in SEM images. The combined interpretation of both metrics therefore improves the robustness of structural assessment and reduces the risk of misinterpretation arising from single-parameter analyses.

A central finding of this study is the strong strain-dependent variability in morphometric responses. In *Nesterenkonia* sp. Act20, fixation time was the dominant determinant of structural variability, with large effect sizes for both Area (ηp² = 0.356) and S/V (ηp² = 0.438), indicating high susceptibility to prolonged chemical processing. In contrast, *Exiguobacterium* sp. S17 exhibited stronger osmium-dependent effects, with significant interactions between fixation duration and OsO_4_ contributing to S/V variability. Osmium tetroxide is known to stabilize lipid-rich domains and reduce membrane damage during processing (Kiernan, 2000; Czerwińska-Główka & Krukiewicz, 2021), suggesting that membrane preservation plays a key role in structural stability in this strain. *Kocuria rosea* CH-021 showed moderate and distributed effects across all factors, consistent with a more robust envelope architecture during fixation and sample preparation.

Collectively, these patterns indicate that fixation response is not a uniform technical artifact but a strain-specific structural response governed by differences in envelope organization, permeability, and structural composition. Rather than reflecting only methodological variability, the observed responses emerge from the interaction between fixative chemistry and intrinsic bacterial architecture, including peptidoglycan organization and surface-associated polymers (Zhu et al., 2021; Whitfield et al., 2022).

Within this general framework, Karnovsky fixation, combined with osmium tetroxide, produced more consistent morphometric outcomes across strains. This effect likely reflects the complementary stabilization of proteins by aldehyde cross-linking and lipid preservation by osmium tetroxide, which together reduce structural loss during processing. However, the impact of osmium tetroxide was not uniform across strains and depended on interactions with fixation time and envelope properties, indicating that the observed morphometric and textural outcomes arise from coupled chemical and biological factors rather than fixative composition alone.

Extremophilic Gram-positive bacteria possess structurally specialized envelopes characterized by thick peptidoglycan layers reinforced by teichoic and lipoteichoic acids, which contribute to mechanical stability, surface charge, and osmotic regulation (Zhu et al., 2021; Whitfield et al., 2022). In addition, stress-adapted membrane composition and cross-linking density may modulate permeability and chemical accessibility (Misra et al., 2013). These features likely contribute to the observed strain-dependent variability in fixation sensitivity by influencing diffusion and dehydration dynamics during sample preparation.

Beyond global geometry, texture analysis using Haralick descriptors provided complementary information about surface ultrastructure. These features, derived from grey-level co-occurrence matrices (GLCM), quantify spatial relationships between pixel intensities and capture nanoscale variations in surface organization (Haralick et al., 1973; Hall-Beyer, 2017). Contrast reflects local intensity variation, energy reflects textural uniformity, entropy quantifies randomness, and homogeneity describes local similarity of grey levels. Although imaging conditions were standardized, part of the observed variability may still arise from topographic, coating-related or acquisition-dependent factors inherent to SEM imaging.

Importantly, several fixation conditions that produced similar morphometric profiles exhibited distinct texture signatures. This dissociation suggests that dimensional stability does not necessarily correspond to identical grayscale texture patterns. Such decoupling may arise from membrane corrugation, partial loss of intracellular components during dehydration, or remodelling of the peptidoglycan network (Gusnard & Kirschner, 1977; Hall-Beyer, 2017). A limitation of the present study is the absence of an independent reference that preserves the native cellular state, such as cryo-SEM, cryo-electron microscopy, or atomic force microscopy of living cells. Consequently, the present analyses allow comparison among fixation protocols but do not permit direct identification of the protocol that most accurately preserves native bacterial ultrastructure. Nevertheless, our findings demonstrate that fixation quality in scanning electron microscopy (SEM) depends on the interaction between chemical processing and strain-specific envelope architecture, and therefore cannot be defined by a single morphological parameter alone, but instead requires a multidimensional analysis based on the integrated evaluation of global cellular geometry and nanoscale surface organization.

## 5. CONCLUSION

Chemical fixation protocols were associated with measurable differences in both morphological and textural properties of extremophilic bacteria in SEM analyses. The absence of a universally optimal protocol highlights the importance of tailoring fixation strategies to strain-specific characteristics. The integration of morphometric and texture-based descriptors provides a quantitative framework for objective evaluation of sample preparation quality. This approach contributes to improving reproducibility and interpretability in ultrastructural studies of microorganisms with specialized cell envelopes.

## Supporting information

MATLAB_Scripts

## 6.#ACKNOWLEDGEMENTS

The authors acknowledge the generous financial support by PIUNT G603 and G703, PICT 2019-3216, CONICET PIP 2023-0040 and USP-T ISYCAV-841/2019 and USP-T ISYCAV-842/2019 projects. VHA is a researcher from the National Research Council (CONICET) in Argentina and a recipient of a Georg Foster Scholarship for Experienced Researchers, Alexander von Humboldt Foundation (2021-2023) and a grant from the Williams Foundation (Fondos Complementarios para Proyectos con Impacto en el Territorio, 2024). FSG was a recipient of a doctoral fellowship from CONICET (2018-2025). FSG is currently an assistant Technician, CPA, CONICET, at CIME. We also thank Luciano Martinez, Hernán Esquivel, Roberto Fanjul and Alejandro Torres for technical assistance. All micrographs were obtained at the Centro Integral de Microscopía Electrónica (CIME), belonging to UNT and CONICET, in Tucumán, Argentina. This manuscript has been released as a Pre-Print at bioRxiv.

## 9. AUTHOR CONTRIBUTIONS

VHA had the original project idea; FSG and VHA designed and carried out the research and wrote the paper; FSG carried out the lab assays and performed the electron microscopy images analysis; VHA provided funding, infrastructure and equipment for the experiments. All authors read and approved the final manuscript.

