## Supplementary material for "FIXATION MATTERS: MULTIDIMENSIONAL EFFECTS OF CHEMICAL PREPARATION ON MORPHOLOGY AND SURFACE TEXTURE OF EXTREMOPHILIC BACTERIA": MATLAB_Scripts: S1_Morphometry_Analysis_S17.m

```
% S1_Morphometry_Analysis_S17.m
% MATLAB script for SEM morphometric analysis
% Developed for quantitative analysis of bacterial ultrastructure
% Associated with:
% "FIXATION MATTERS: MULTIDIMENSIONAL EFFECTS OF CHEMICAL
% PREPARATION ON MORPHOLOGY AND SURFACE TEXTURE OF
% EXTREMOPHILIC BACTERIA"
%
% Input:
%   Dataset_Supplementary.xlsx
%   Sheet: "Morphometric dataset_S17"
%
% Output:
%   Statistical analysis tables
%   Post hoc comparisons
%   Morphometric plots
%
% Software:
%   MATLAB® (The MathWorks Inc., Natick, MA, USA)

function S1_Morphometry_Analysis_S17()

    % Load dataset
    archivo = 'S3_Morphometric_and_Texture_Datasets.xlsx';
    hoja = 'Morphometric dataset_S17';

    datos = readtable(archivo, 'Sheet', hoja);

    % Remove rows with missing values
    datos = rmmissing(datos);

    % Create output folders
    carpeta_graficos = 'Morphometry_Results';
    carpeta_anova = 'ANOVA_Results';

    if ~exist(carpeta_graficos, 'dir')
        mkdir(carpeta_graficos);
    end

    if ~exist(carpeta_anova, 'dir')
        mkdir(carpeta_anova);
    end

    % Response variables
    variables = {'Area', 'SV'};

    for v = 1:length(variables)

        var = variables{v};

        fprintf('\nANALYSIS FOR VARIABLE: %s\n', var);

        % Select required columns
        tbl_tmp = datos(:, {'Fixative', 'Time', 'OsO4', var});
        tbl_tmp = rmmissing(tbl_tmp);

        % Three-way ANOVA
        [~, tbl_stats, stats, ~] = anovan(tbl_tmp.(var), ...
            {tbl_tmp.Fixative, tbl_tmp.Time, tbl_tmp.OsO4}, ...
            'model', 'full', ...
            'varnames', {'Fixative', 'Time', 'OsO4'}, ...
            'display', 'on');

        % Multiple comparisons for triple interaction
        [c, m, ~, gnames] = multcompare(stats, ...
            'Dimension', [1 2 3], ...
            'Display', 'off');

        letras_inter = calculaLetrasSignificanciaMC(c, gnames);

        % Save post hoc letters, means and SEM
        grupo_tabla = table(gnames, m(:,1), m(:,2), letras_inter, ...
            'VariableNames', {'Group', 'Mean', 'SEM', 'Letter'});

        writetable(grupo_tabla, ...
            fullfile(carpeta_graficos, ...
            ['posthoc_letters_' var '.xlsx']));

        % Bar plot with significance letters
        figure('Name', ['Barplot - ' var], 'Visible', 'off');

        hold on

        for i = 1:length(m(:,1))

            bar(i, m(i,1), 'FaceColor', [0.2 0.6 0.8]);

            errorbar(i, m(i,1), m(i,2), '.k', ...
                'LineWidth', 1.5);

            text(i, ...
                m(i,1) + m(i,2) + 0.05*max(m(:,1)), ...
                letras_inter{i}, ...
                'HorizontalAlignment', 'center', ...
                'FontSize', 12);
        end

        title(['Mean ± SEM - ' var]);

        ylabel(var);

        xticks(1:length(gnames));
        xticklabels(gnames);

        xtickangle(45);

        grid on;
        hold off;

        saveas(gcf, ...
            fullfile(carpeta_graficos, ...
            ['barplot_' var '.png']));

        % Boxplot with mean markers
        figure('Name', ['Boxplot - ' var], 'Visible', 'off');

        grupo_combinado = strcat( ...
            string(tbl_tmp.Fixative), '-', ...
            string(tbl_tmp.Time), '-', ...
            string(tbl_tmp.OsO4));

        boxplot(tbl_tmp.(var), grupo_combinado);

        title(['Boxplot - ' var]);

        ylabel(var);

        xtickangle(45);

        grid on;

        % Add mean markers
        hold on

        grupos = unique(grupo_combinado, 'stable');

        for i = 1:length(grupos)

            idx = grupo_combinado == grupos(i);

            media = mean(tbl_tmp.(var)(idx));

            plot(i, media, 'o', ...
                'MarkerSize', 3, ...
                'MarkerEdgeColor', 'r', ...
                'MarkerFaceColor', 'none');
        end

        hold off

        saveas(gcf, ...
            fullfile(carpeta_graficos, ...
            ['boxplot_' var '.png']));

        % Export ANOVA table
        tabla_anova = cell2table(tbl_stats(2:end,:), ...
            'VariableNames', ...
            {'Source', 'SumSquares', 'DF', ...
            'Singular', 'MS', 'F', 'p'});

        writetable(tabla_anova, ...
            fullfile(carpeta_anova, ...
            ['anova_' var '.xlsx']));

        % Export means and SEM by group
        medias_sem = grpstats(tbl_tmp, ...
            {'Fixative', 'Time', 'OsO4'}, ...
            {'mean','sem'}, ...
            'DataVars', var);

        writetable(medias_sem, ...
            fullfile(carpeta_anova, ...
            ['means_sem_' var '.xlsx']));
    end
end

function letras = calculaLetrasSignificanciaMC(c, gnames)

    % Calculate significance letters from multcompare results

    n = length(gnames);

    sigMat = true(n);

    for i = 1:size(c,1)

        g1 = c(i,1);
        g2 = c(i,2);
        pval = c(i,6);

        if pval < 0.05

            sigMat(g1,g2) = false;
            sigMat(g2,g1) = false;
        end
    end

    alf = 'abcdefghijklmnopqrstuvwxyz';

    groupLetters = repmat({''}, n, 1);

    for letter = 1:length(alf)

        for i = 1:n

            if groupLetters{i} == ""

                puedeAsignar = true;

                for j = 1:n

                    if j ~= i && ~sigMat(i,j)

                        if ~isempty(groupLetters{j}) && ...
                                contains(groupLetters{j}, alf(letter))

                            puedeAsignar = false;
                            break
                        end
                    end
                end

                if puedeAsignar

                    groupLetters{i} = ...
                        [groupLetters{i}, alf(letter)];
                end
            end
        end

        if all(cellfun(@(x) ~isempty(x), groupLetters))
            break
        end
    end

    letras = groupLetters;
end
```
